# Offset analgesia and onset hyperalgesia: A comparison between women with non-suicidal self-injury and healthy women

**DOI:** 10.1101/2025.02.07.637056

**Authors:** Jens Fust, Granit Kastrati, Johan Bjureberg, Nitya Jayaram-Lindström, Clara Hellner, Eva Kosek, Peter Fransson, Karin B. Jensen, Maria Lalouni

## Abstract

**Introduction:** Individuals who engage in non-suicidal self-injury (NSSI) exhibit reduced pain sensitivity compared to the general population. It has been argued that this hypoalgesic characteristic may be attributable to hyper-effective pain modulation. However, empirical support for this theory remains inconsistent.

**Objective:** The aim of the study was to use a combined offset analgesia (OA) and onset hyperalgesia (OH) protocol to investigate if women with NSSI have a propensity to inhibit nociceptive signals to a higher degree (stronger OA response) and facilitate nociceptive signals to a lesser degree (weaker OH response), compared to a control group.

**Methods:** Data was collected from 76 women, 18-35 age (37 with NSSI and 39 healthy controls). The OA and OH protocol was combined with functional magnetic resonance imaging (fMRI).

**Results:** The NSSI group displayed a weaker OH response, compared to the control group. This suggests that women with NSSI do not facilitate nociceptive signals to the same extent as healthy women. However, there were no significant differences between the groups regarding OA. Across all participants, we observed stronger activation in the primary sensory cortex during the OH condition, compared to the control condition.

**Conclusions:** The results offer partial support for the general hypothesis that women with NSSI demonstrate enhanced pain modulation.

## Introduction

Individuals who engage in non-suicidal self-injury (NSSI) tend to be less sensitive to pain than the general population. Not only do many individuals with NSSI report feeling little or no pain while harming themselves (Izadi-Mazidi et al., 2019; Lloyd-Richardson et al., 2007; Zetterqvist et al., 2013), but also, in laboratory studies the NSSI population display higher pain threshold, higher pain tolerance, and report lower pain intensity when being exposed to a painful stimulus, compared to control groups (Koenig et al., 2016). Longitudinal data suggest that high pain tolerance is a risk factor for severe NSSI engagement (Boyne & Hamza, 2024).

It has been theorized that this hypoalgesic trait could be explained by hyper-effective pain modulation (Defrin et al., 2020; Lalouni et al., 2022), i.e. a propensity to inhibit nociceptive signals to a higher degree and facilitate nociceptive signals to a lesser degree. However, the evidence for this theory is mixed. In our previous study (Defrin et al., 2020; Lalouni et al., 2022) women with NSSI displayed greater conditioned pain modulation response, a measure of endogenous pain inhibition, compared to the control group. In contrast, Leone et al. (Leone et al., 2021) reported that adolescents with NSSI demonstrated weaker conditioned pain modulation response than a control group. Moreover, NSSI has not been associated with abnormal temporal summation of pain, which is believed to be a measure of central aspects of pain facilitation (Lalouni et al., 2022). At this time, considering the apparently inconsistent results, it is hard to draw any strong conclusions regarding the pain modulatory system of the NSSI population.

Offset analgesia (OA) is an alternative protocol to characterize pain inhibition (Grill & Coghill, 2002). While conditioned pain modulation could be said to measure spatial filtering of nociceptive information, OA is related to temporal filtering of nociceptive information (Nahman-Averbuch et al., 2014). The OA protocol compares a constant noxious heat stimulation with an identical heat stimulation, apart from a brief period of increased temperature. The short deviation in temperature leads to a disproportionate reduction of pain intensity, compared to the constant heat stimulation. It has also been demonstrated that a hyperalgesic response can be produced using an inverted version of the OA protocol, called onset hyperalgesia (OH) (Alter et al., 2020). The physiological basis of OA is still largely unknown. Although centrally acting drugs seem not to affect the OA response (Boye Larsen et al., 2022), there are studies suggesting involvement of the central as well as the peripheral nervous system (Ligato et al., 2018; Sitsen et al., 2020; Sprenger et al., 2018). Brain imaging studies have found that the OA response is associated with activity in brainstem (Derbyshire & Osborn, 2009; Nahman-Averbuch et al., 2014; Yelle et al., 2009; Zhang et al., 2018) and dorsolateral prefrontal cortex (Alter et al., 2022; Nahman-Averbuch et al., 2014; Yelle et al., 2009; Zhang et al., 2018).

Because of the mixed results in the studies investigating the role of pain modulation in NSSI-related hypoalgesia, there is a need to further characterize the pain modulation of the NSSI population. In the present study, participants with ongoing NSSI behavior and age-matched healthy controls conducted a combined OA and OH protocol. The advantage of the combined protocol is that, through a minor adjustment of the thermal stimulation, it allows for the assessment of both endogenous pain inhibition and facilitation. Our hypotheses were:

1. The NSSI group will display a stronger OA response, compared to healthy individuals
2. The NSSI group will display a weaker OH response, compared to healthy individuals

To our knowledge, this is the first study of OH in a clinical population, and the first study combining OH with brain imaging.

## Methods

### Participants

Our power analysis was aimed at detecting differences in pain threshold between NSSI participants and controls. Based on a meta-analysis by Koenig et al. (Koenig et al., 2016), we estimated that 29 participants in each group were needed to achieve 80% power. To ensure the ability to detect smaller effects and account for potential incomplete data, we recruited 40 participants for the NSSI group and 40 participants for the control group. We decided to only recruit female participants because NSSI is more common in women, and we did not think we would be able to recruit the number of male participants to investigate sex differences. fMRI data was collected from 34 participants in the NSSI group. Of the six participants for whom fMRI data was not collected: one participant could not remove their piercing; one could not remove their jewelry; one aborted the scan due to claustrophobia; two experienced MR scanner malfunctions; and in one case, the experimenters aborted the scan because the rated pain intensity was deemed too high. Three of these six participants completed the experiment outside the scanner, allowing us to collect their pain ratings. As a result, we collected pain ratings from 37 participants in the NSSI group. We collected fMRI data and pain ratings from 39 participants in the HC group. One participant could not complete the experiment because of technical difficulties with the thermal stimulator. A description of the characteristics of the NSSI group and the control group can be found in a previously published study using the same participants (Lalouni et al., 2022).

### Recruitment

Participants with NSSI and controls were recruited between April 2019 and June 2020, through flyers in waiting rooms at outpatient psychiatric clinics (to recruit NSSI participants) and advertisements in social media (NSSI participants and controls). The advertisement for controls was adjusted during the inclusion phase to match the age and educational level of the NSSI participants. General inclusion criteria were: a) woman, b) 18-35 years, and c) right-handed. General exclusion criteria were: d) chronic inflammatory, autoimmune, or other somatic disorder requiring treatment, e) pain condition, f) contraindication for fMRI (e.g., metal implant, pregnancy, claustrophobia), g) suicide attempts during the last year, h) suicidal plans or acute risk for suicide. Specific inclusion criteria for participants with NSSI: i) engaged in self-injury on ≥ 5 days within the past year. Specific exclusion criteria for controls: j) treatment for depression or anxiety. Participants with NSSI received a remuneration of approximately $158 (1500 SEK) and controls received approximately $126 (1200 SEK). Participants with NSSI received the higher amount because their participation included a visit to a clinical psychologist (JB) to confirm current NSSI behavior and to assess the risk of participating in the study. The study was approved by the Regional Ethical Review Board in Stockholm (2018/1367-31/1).

### Setting

Data was collected at the MR center at Karolinska University Hospital in Stockholm, Sweden, between May 2019 and August 2020. The study was part of a larger research project, assessing the relationship between NSSI and pain modulation.

### Thermal stimulation and pain assessment

Heat stimuli were administered with a thermal stimulator (Somedic Senselab AB, Hörby, Sverige) with a 30 × 30 mm thermal probe. Temperature increased and decreased at a rate of 5°C/s. Participants used a trackball to rate their pain intensity on a numeric rating scale (NRS) that was displayed on a screen, marked with integers ranging from 0 to 10 on a horizontal line. Participants were instructed verbally that 0 represented “no pain” and 10 “worst imaginable pain”.

### Functional MRI acquisition

Magnetic resonance images were collected with a 3T General Electric 750 MR scanner. We acquired functional scans using T2*-weighted single-shot gradient echo planar imaging with the following parameters: repetition time/echo time = 2000/30 ms, flip angle = 70°, field of view = 220 × 220 mm, matrix size = 72 × 72, 42 slices, slice thickness = 3 mm with a 0.5-mm gap, axial acquisition plane, interleaved slice acquisition. Anatomical images were collected with a high-resolution BRAVO 3D T1-weighted image sequence (1 × 1 × 1-mm voxel size, 176 slices). A total number of 220 fMRI volumes were acquired in the experiment.

### Pain calibration

Pain calibration of the thermal stimulation was conducted on a visit before the experiment. The thermal probe was attached to participants’ left calf. Each thermal stimulation lasted 5 seconds, with an interstimulus interval of 35 seconds. Participants used a handheld trackball to rate their pain intensity on a 0–10 NRS that was displayed on a screen after each stimulation. First, starting from 38°C, temperature was raised 1°C for each subsequent stimulation, until the participants rated their pain above NRS 1/10. Second, participants were administered random temperatures, based on their pain threshold. The participants were exposed to between 13–24 heat stimulations (depending on the pain sensitivity of the participant). The temperature never exceeded 50°C, to avoid the risk of tissue damage. We applied linear regression (temperature ∼ pain rating) to the thermal stimulations with random temperatures to predict the temperature of each participant’s pain rating of NRS 5/10.

### Experimental procedure

The thermode was attached to the participants’ left calf while the participants were lying inside the MR scanner. Before the start of the experiment, the participants were asked to continuously rate 15 seconds of thermal stimulation twice. The reason was to confirm the individually calibrated temperature and also familiarize the participants with the rating procedure. The first thermal stimulation was 2°C below the calibrated temperature and the second thermal stimulation was the calibrated temperature. If the maximum pain rating of the calibrated temperature was between NRS 4/10 and 6/10, this temperature was used as the “medium temperature” in the experiment, otherwise the procedure was repeated with a higher or lower temperature until the desired pain rating was reached. The OA and OH protocol can be divided up in 3 time intervals: T1, T2, and T3 (see Fig. 1 for a visual representation). During T1 (0–6.5 seconds), the temperature of the thermode increased from a non-painful temperature (38°C) to the individual calibrated medium temperature. T1 continued approximately 5 seconds after the thermode reached the medium temperature. During T2 (6.5–12 seconds), the temperature was either kept stable (control), increased 2°C (OA), or decreased 2°C (OH) from the medium temperature. T2 continued for 5 seconds after the thermode reached the assigned temperature. During T3 (12–33 seconds), the thermode returned to the medium temperature, and after approximately 20 seconds, the thermode returned to the non-painful baseline temperature. Pain ratings were continuously registered until approximately 10 seconds after the thermode returned to the baseline temperature. The three conditions (OA, OH, and control) were repeated twice. The order of the conditions was randomized for each participant. Every condition was followed by a 50 second inter-trial interval where the participants were exposed to the baseline temperature of 38°C. The thermode was not moved between trials.

**Figure 1:**
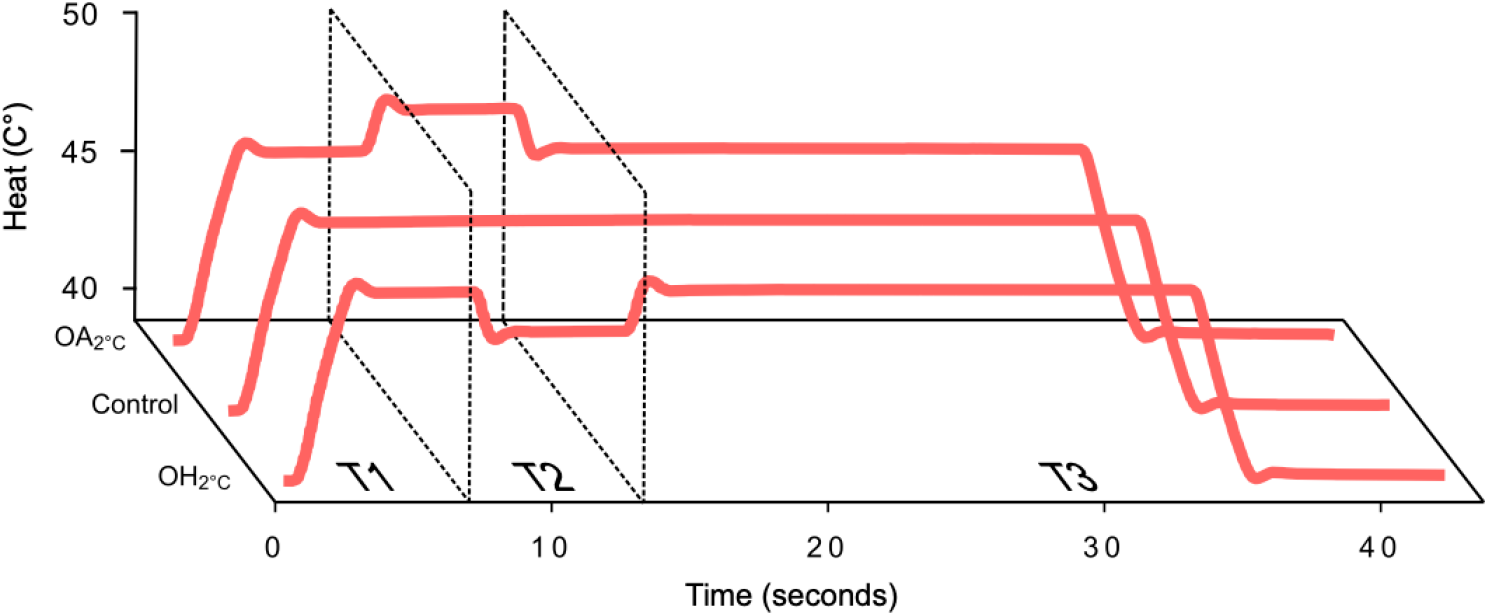
All conditions.

### Data processing and statistical analysis

The analysis plan was registered on Open Science Framework on October 25th 2021 (https://osf.io/3bquj), after data was collected but before data was analyzed.

### Pain ratings

First, individual pain ratings were downsampled to 10 Hz in order to adjust for occasional frame drops (pain ratings were originally collected for each frame rate for the computer screen). Second, we calculated the mean pain rating during the last 12 seconds of heat stimulation (T3 interval, between 21-33 seconds of stimulation), for each condition and each participant (Fig. 2). This 12 second interval was decided based on the results of our previous experiments (Fust et al., 2020). Third, we created two mixed effect models using the lme4 and lmerTest packages in R. One model estimated the OA response and compared the OA response between groups. The other model estimated the OH response and compared the OA response between groups. In both models, condition, group and the interaction between group and condition was specified as fixed effects. The models also included a by-subject random effect for condition.

**Figure 2:**
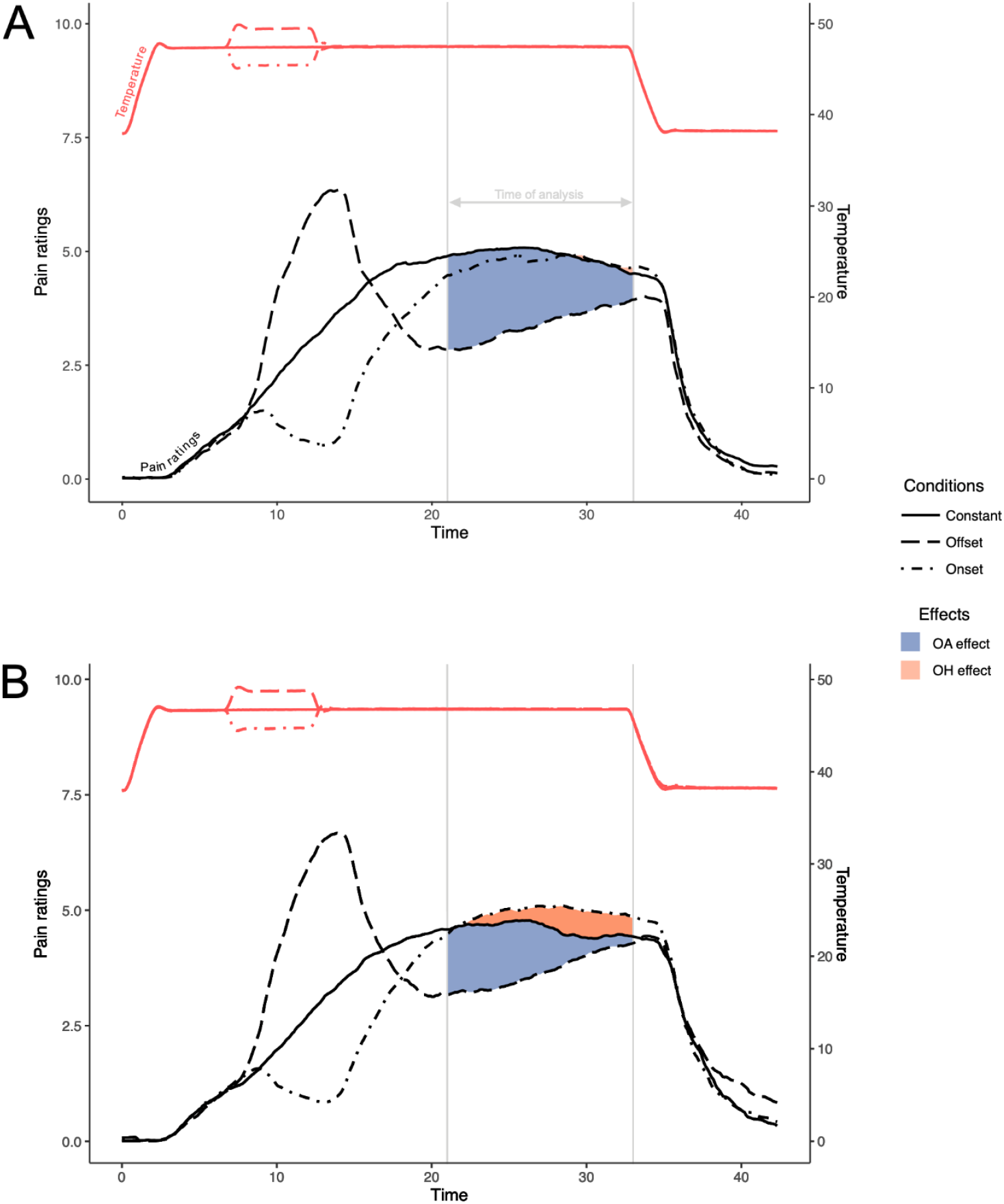
Mean pain ratings and temperature (degrees Celsius) for the NSSI group (A) and HC group (B). OA, offset analgesia; OH, onset hyperalgesia; NSSI, Non-suicidal self-injury; HC, healthy control.

We also calculated the mean OA and OH response for each participant to test the correlation of the effects. To avoid comparing dependent measures of the OA and OH response, the mean OA response was defined as the difference between the mean of the pain ratings in the two OA conditions and the pain ratings in one of the two control conditions (during the last 12 seconds of heat stimulation), and the mean OH response was defined as the difference between two OH conditions and the other control condition. If the OA condition was first in the randomized sequence, the OA conditions were paired with the first control condition, and the OH conditions were paired with the second control condition in the randomized sequence, and vice versa.

### fMRI

Quality control of the imaging data was conducted using MRIQC (https://mriqc.readthedocs.io/en/stable/). Preprocessing was performed with the fMRIPrep pipeline (https://fmriprep.org/), incorporating realignment and spatial normalization to the Montreal Neurological Institute (MNI) standard space. Functional images were subsequently smoothed using an 8-mm full-width at half-maximum (FWHM) Gaussian kernel. Statistical analyses were conducted in SPM12 within MATLAB 2014 (The MathWorks, Inc., Natick, MA). Participants with excessive head movement during the fMRI scan were removed if they had more than 20% of scans with more than 0.5 in framewise displacement during the entire experiment. Two controls and two NSSI participants were removed from the final analysis based on this predefined cutoff.

The fMRI BOLD signal for the OA condition across groups was determined on the individual level (first-level) by subtraction analysis: [OA - Control]. A group analysis (second-level) of the OA main effect [OA - Control] across groups was performed using a one-sample t-test. Next, the interaction group/condition was performed: NSSI [OA - Control] > Healthy controls [OA - Control] using a two-sample t-test, and the opposite: NSSI [OA - Control] < Healthy controls [OA - Control]. A priori brain regions for region of interest (ROI) analyses for OA include: Brainstem, dorsolateral prefrontal cortex (dlPFC), rostral Anterior Cingulate Cortex (rACC), and thalamus. In addition to the ROI results, we also report whole-brain results for the OA effect. All group analyses were performed using an initial statistical threshold of p<.001, uncorrected for multiple comparisons, and an extent threshold of 20 voxels. Activation clusters were considered significant at p<.05, Family Wise Error-corrected for multiple comparisons.

The fMRI BOLD signal for the OH condition across groups was determined on the individual level (first-level) by subtraction analysis: [OH - Control]. A group analysis (second-level) of the OH main effect [OH - Control] across groups was performed using a one-sample t-test. Next, the interaction group/condition was performed: NSSI [OH - Control] > Healthy controls [OH - Control] using a two-sample t-test, and the opposite: NSSI [OH - Control] < Healthy controls [OH - Control]. A priori brain regions for ROI analyses for OH include: S1 and S2, insula, Anterior Cingulate Cortex (ACC), and inferior frontal cortex (IFC). In addition to the ROI results, we also report whole-brain results for the OH effect. All group analyses were performed using an initial statistical threshold of p<.001, uncorrected for multiple comparisons, and an extent threshold of 20 voxels. Activation clusters were considered significant at p<.05, Family Wise Error-corrected for multiple comparisons. All ROIs were based on the Automated anatomical labeling (AAL) atlas 3, with the exception for the brainstem, which was based on meta-analysis (Linnman et al., 2012).

## Results

### Pain ratings

#### Offset analgesia

Model estimated OA response across groups was NRS -0.90 (95% CI -1.43, -0.36, P = 0.001; Fig. 3). The model estimated difference in OA response between the NSSI group and control group was not statistically significant (NRS -0.63, 95% CI -1.39–0.13, P = 0.10; Fig. 3).

**Figure 3:**
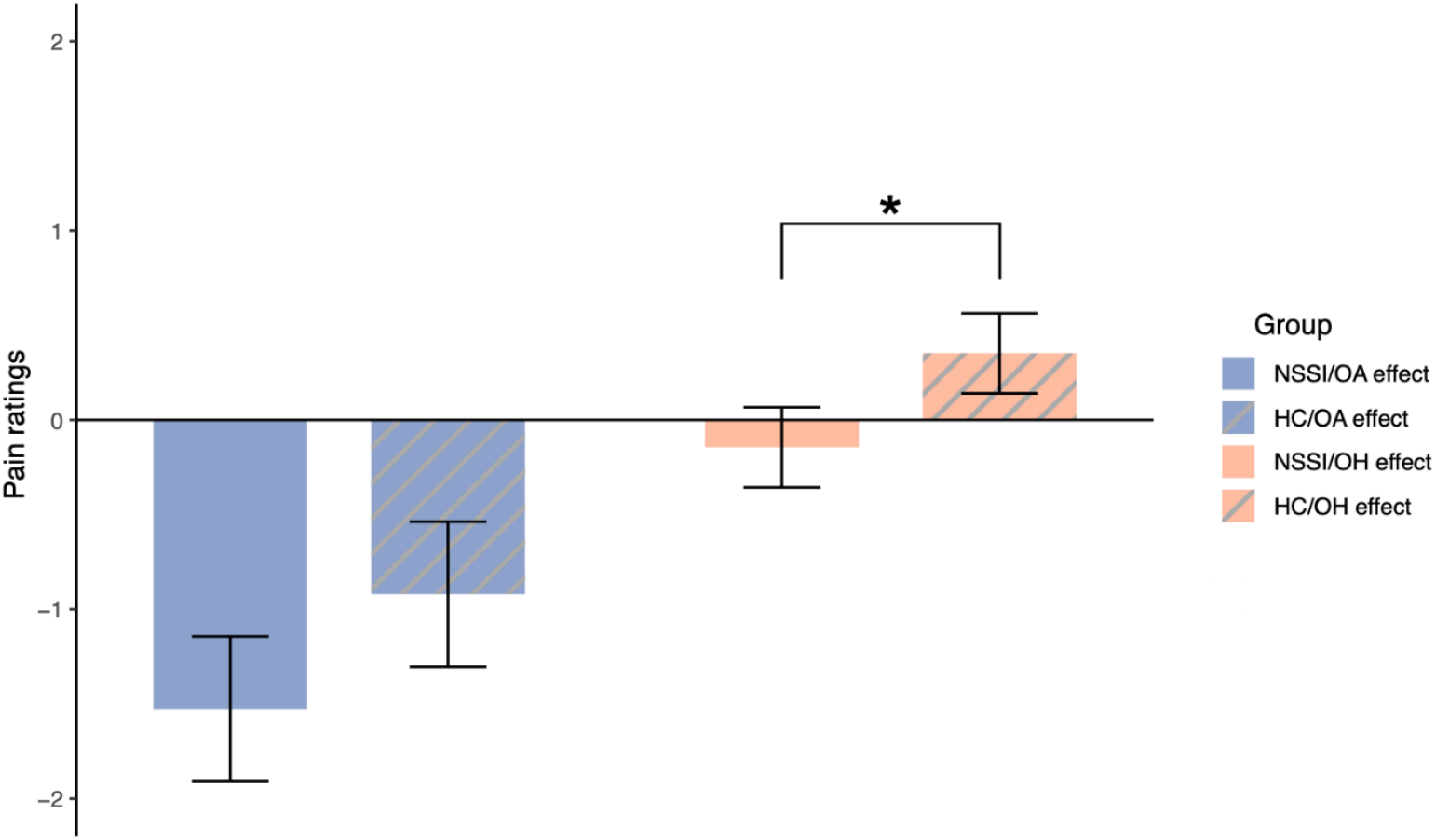
Mean OA and OH effects for the NSSI and HC group. Error bars represent ±1 within-subject standard error of the mean. OA, offset analgesia; OH, onset hyperalgesia; NSSI, Non-suicidal self-injury; HC, healthy control.

#### Onset hyperalgesia

Model estimated OH response across groups was NRS 0.38 (95% CI 0.10–0.66, P < 0.001; Fig. 3). The model estimated difference in OH response between the NSSI group and control group was statistically significant (NRS -0.54, 95% CI -0.13, -0.93, P < 0.001; Fig. 3). The direction of the result indicates a weaker OH effect in the NSSI group compared to controls.

#### Correlation between OA response and OH response

There was a significant correlation between OA response and OH response (r = 0.28; P = 0.01) across groups.

### fMRI

#### Offset analgesia

There were no significant activation clusters related to the OA response across groups or between groups, neither in the ROIs or whole-brain analyses.

#### Onset hyperalgesia

During OH, there was a significant increase of brain activity in the primary somatosensory cortex across groups (Fig. 4), approximately corresponding to the contralateral leg area vis a vis the left-sided pain stimulation on the calf (188 voxels, pFWE = 0.032, MNI coordinate for cluster peak voxel x = 14, y = –37, z = 68). The results suggest an increased activation of the cortical representation of the stimulated leg area during OH. There were no significant clusters between groups, neither for the ROIs or whole-brain analyses. For exploratory reasons, a lowered initial threshold of p<.05 uncorrected of the whole brain indicates that healthy controls had more increases of brain activations during OH, compared to NSSI, represented in primary and secondary somatosensory cortex. Yet, that observation needs to be formally tested in future studies.

**Figure 4:**
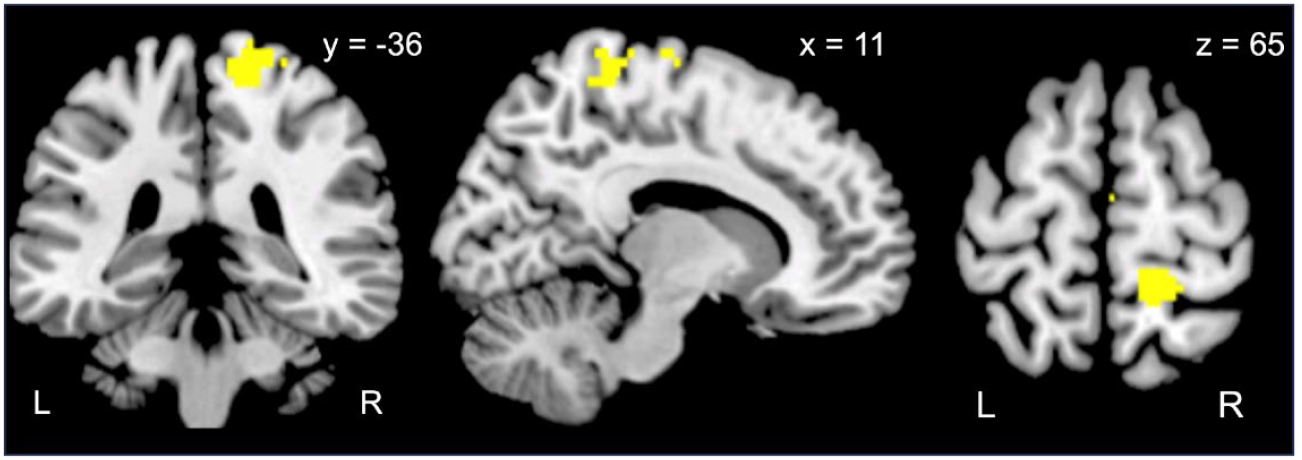
Representation of fMRI BOLD signal associated with the OH response in the primary sensory cortex, compared to the control condition. The cluster in yellow is a binary representation of the significant activation contralateral to the pain stimulation. The location of the cluster is indicated by MNI (x, y, z) coordinates. BOLD, blood-oxygen-level-dependent; OH, onset hyperalgesia.

## Discussion

In the present study, women with ongoing NSSI behavior and age-matched control group were tested with a combined OA and OH protocol, during fMRI. Two hypotheses were tested: the NSSI group will display (H1) stronger OA response and (H2) weaker OH response, compared to the control group. The study was part of a larger research project, studying the relationship between NSSI and pain modulation.

In line with the second hypothesis, the women with NSSI behavior displayed weaker OH responses compared to the control group. This suggests that women with NSSI do not facilitate nociceptive signals to the same extent as women without NSSI behavior, which supports the idea that women with NSSI have a hyper-effective pain modulation system. The mean difference in pain ratings between groups was small but this was probably due to the low average OH response in this study.

In contrast to the first hypothesis, the OA response was not significantly higher in the NSSI group compared to the control group. The result does not give further support to the notion that the NSSI population has more effective inhibition of pain suggested by some of the earlier studies using conditioned pain modulation, including our own study (Lalouni et al., 2022. Although, it is not clear how other measures of central inhibition of pain, such as conditioned pain modulation, is related to OA. Based on previous studies it is likely that OA does not strongly correlate with conditioned pain modulation {Nahman-Averbuch, 2014 #23). It is possible that conditioned pain modulation is more associated with the hypoalgesic trait that has been observed in the NSSI population than OA.

Across all participants, we registered stronger activation in a region in the primary sensory cortex during the OH condition, compared to the control condition. This was not due to differences in temperature because the temperature was identical in both the OH and control condition during the time of analysis. However, it is still possible that the BOLD signal is related to the afferent nociceptive signal. The nociceptive signal may be upregulated at the spinal and/or peripheral level before reaching the brain. Regarding OA, it has been found that the OA response is associated with downregulation of the nociceptive signal in the spinal cord {Sprenger, 2018 #16}. In the future it would be a good idea to study OH in combination with fMRI without concurrent pain ratings. Even though a null result is technically considered as inconclusive, it is surprising that we did not find any significant clusters associated with the OA response. Previous studies, including about a quarter to a third of the sample size of this study, reported OA-related findings in the brainstem (Derbyshire & Osborn, 2009; Nahman-Averbuch et al., 2014; Yelle et al., 2009) and dorsolateral cortex (Alter et al., 2022; Nahman-Averbuch et al., 2014; Yelle et al., 2009). There is a need for a large-scale study on the neural correlates of OA in order to estimate the true effects on pain inhibitory circuitry in the brain.

We found a correlation between OA and OH responses, across all of the participants. A stronger OA response was associated with a weaker OH response, and vice versa. In line with earlier studies (Alter et al., 2020; Fust et al., 2020), this correlation was weak. The weak correlation suggests that OA and OH responses could be related to different pain modulatory mechanisms. Future studies should aim to disentangle the psychophysical properties and biological correlates of OA and OH. For example, it is possible that the OH response is more susceptible to pharmacological manipulations than the OA response.

A limitation of this study is that we compared women with NSSI with a non-psychiatric control group. This makes it hard to determine if the difference between the groups is attributed to NSSI or other psychiatric traits in the NSSI group. There was only a small difference in OH response between the groups. It is possible that this group difference could be affected by other non-pain related factors such as attention or motivation. Another limitation is the sample size of the study, which did not allow us to detect subtle differences between the groups, which could be relevant. This is especially true regarding the fMRI results, because effect sizes in fMRI research are in general small (Poldrack et al., 2017). Finally, this study used a slightly different OH protocol (Fust et al., 2020) than the original OH study by Alter et al. (Alter et al., 2020). The original OH protocol seems to produce stronger and more reliable OH responses and should be used in future studies of OH.

Observational data support the notion that increased pain tolerance constitutes a risk factor for severe NSSI engagement, which may be explained by the role of pain as a barrier to self-harm (Boyne & Hamza, 2024). If it can be established that individuals who engage in NSSI behavior exhibit altered pain modulation, this could at least partially account for the elevated pain tolerance observed in this population. Evidence of hyper-effective pain modulation in the NSSI would also provide a potential target for pharmacological or behavioral interventions.

To our knowledge, this is the first study using OH to investigate pain modulation in a psychiatric patient group. To assess whether OH is a clinically relevant measure of endogenous pain modulation, it would be valuable to compare a patient group with a pronociceptive pain modulation profile, such as patients with nociplastic pain with a group of healthy individuals. The interaction between pain modulation in the NSSI population and patients with nociplastic pain, might be key to understanding how individuals move along the nociceptive spectrum.

## Acknowledgements

This work was supported by grants from the Swedish Research Council (2019-02936), StratNeuro Collaborative Grant, and a donation from Leif Lundblad to pain research. Open access funding provided by Karolinska Institute. Data supporting this study cannot be made openly available due to legal reasons but will be made available upon reasonable request. Analysis scripts can be found at Open Science Framework: https://osf.io/x8hjk.

